# Cafe: An integrated platform for exploring cell fate

**DOI:** 10.1101/2025.02.04.636565

**Authors:** Zhaoyang Huang, Haonan Ma, Yuchuan Peng, Chenguang Zhao, Liang Yu

**Affiliations:** School of Computer Science and Technology, Xidian University, Xi’an 710071, Shaanxi, China; Department of Rehabilitation Medicine, Xijing Hospital, Fourth Military Medical University, Xi’an, Shaanxi, China

**Keywords:** Cell fate prediction, Trajectory inference, Benchmark

## Abstract

Cell fate prediction from single-cell omics data is a central task for reconstructing developmental dynamics and inferring lineage relationships, but existing tools remain fragmented and often tailored to specific trajectory structures. Here we present Cafe (CelluAr Fate Explorer), an integrated platform for cell fate analysis that unifies trajectory-aware data storage, heterogeneous inference methods, benchmarking, visualization, and downstream interpretation. Cafe standardizes diverse inputs into a common milestone-network representation, supports both native and Dynverse-compatible methods, and evaluates performance across representative benchmark criteria. Applied to representative case studies across atlas-scale and lineage-resolved datasets, Cafe recovers coherent developmental trajectories and provides benchmark-guided method selection across data types and biological contexts. A browser-based interface built on cellxgene enables interactive trajectory inspection. Cafe also supports downstream analyses, including driver-gene prioritization and gene-regulatory-network inference. As an open, community-driven platform, Cafe is designed to evolve continuously with contributions from developers and users, expanding method coverage and functionality for cell fate prediction.

## Introduction

Single-cell sequencing technologies, particularly single-cell RNA sequencing (scRNA-seq), have transformed the study of dynamic biological processes by enabling transcriptome profiling at single-cell resolution. These data provide a powerful basis for reconstructing developmental trajectories and inferring cell fate decisions during differentiation, organogenesis, and disease progression^[1-10]^. Beyond clustering and annotation, trajectory inference has become a central computational task for ordering cells along developmental paths and recovering lineage relationships from static snapshots of heterogeneous populations. Over the past decade, a wide range of trajectory inference methods has been proposed, including pseudotime-based approaches such as Palantir^[11]^ and Cytotrace^[12]^, graph-based methods such as Monocle^[11]^, and RNA-velocity-based models such as scVelo and Dynamo^[13, 14]^. Because these methods are often optimized for different trajectory structures, data modalities, and prior assumptions, selecting an appropriate method for a given dataset remains a nontrivial challenge.

To address this problem, several benchmarking frameworks have been developed to compare trajectory inference methods under standardized settings. Dynverse established an important foundation by evaluating a large collection of methods across many datasets^[15]^, but its method coverage has not kept pace with recent developments, particularly RNA-velocity-based and other emerging approaches^[16, 17]^. Moreover, trajectory analysis is not only a question of algorithmic accuracy, but also of data preparation, interoperability, downstream interpretation, and usability. In practice, reseachers still face fragmented workflows, inconsistent data formats, and substantial overhead in switching between methods, environments, and visualization tools^[18]^.

Cafe (CelluAr Fate Explorer) was developed to address these limitations by providing an integrated and extensible platform for cell fate analysis. Inspired by general-purpose single-cell ecosystems such as Scanpy^[19]^, Scverse^[20]^, OmicVerse^[21]^, and Pertpy^[22]^,Cafe focuses specifically on trajectory inference and cell fate prediction. Its design combines four major components: the data module standardizes single-cell and benchmark datasets through a trajectory-aware data structure that preserves both expression information and reference lineage annotations; the method module provides a unified interface for a broad collection of trajectory inference methods, including native Cafe implementations and Dyn-verse-compatible wrappers; the visualization module visualize the abundant trajectory results among various methods by static image and dynamic interactive visualization; the explorer module focus on benchmark and downstream analyses such as driver-gene discovery and enrichment analysis. Subsequent results show that this framework standardizes heterogeneous datasets and trajectory outputs while supporting native and Dynverse-compatible methods across different trajectory structures, reconstructs developmental trajectories in atlas-scale gastrulation data through a divide-and-conquer strategy, and provides a cellxgene-based interactive analysis environment for trajectory inspection and downstream exploration. Taken together, Cafe provides a practical foundation for systematic trajectory analysis and sets the stage for future expansion toward atlas-scale, multi-modal, and more interactive cell fate exploration.

## Result

### Cafe provide comprehensive analysis tools for cell fate prediction

Cafe provides an end-to-end framework for cell fate analysis that integrates data management, trajectory inference, visualization, exploration within a single workflow, as shown in as illustrated in Figure 1.

**Figure 1.**
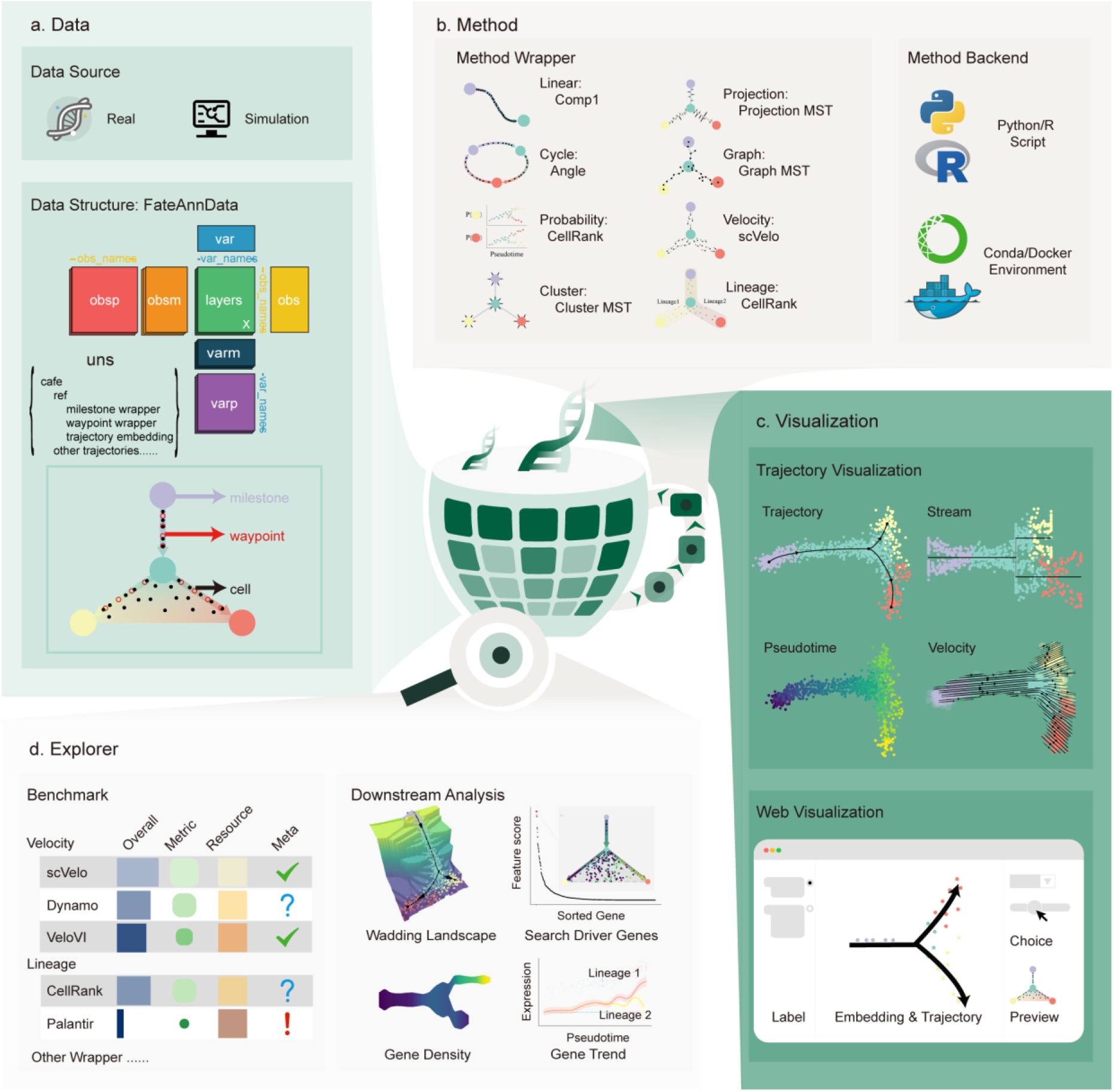
Framework of Cafe. **a**, the data module organizes real and simulated datasets by FateAnnData data structure with trajectory-aware annotation. **b**, the method module standardizes diverse trajectory outputs into a unified milestone-network representation and supports Python/R base on conda/docker software environment. **c**, the visualization module provides multiple views such as trajectory, stream, pseudotime, velocity and so on for trajectory visualization, together with an interactive web based on cellxgene. **d**, the explorer module combines benchmark comparison, metadata reporting, and downstream analysis for biological interpretation, including Waddington landscapes, driver-gene ranking, gene-density, and gene-trend analyses.

The data module is centered on FateAnnData, an AnnData-compatible container that stores both real and simulated datasets together with trajectory-aware annotations, including milestone and waypoint information. This design preserves compatibility with the scverse ecosystem while retaining the topological context required for trajectory reconstruction and downstream interpretation^[20]^. The Method module standardizes heterogeneous outputs through a unified wrapper system and supports Python or R scripts as well as conda/docker-based execution. By converting diverse method-specific outputs into a common milestone-network representation, it decouples algorithm implementation from the user environment and enables direct comparison across linear, cyclic, probability-based, cluster-based, projection-based, graph-based, velocity-based, and lineage-based methods. The visualization module provides coordinated views for trajectory, stream, pseudotime, and velocity rendering, and its interactive web interface allows users to select labels, inspect embeddings and trajectories jointly, and preview candidate lineages in real time. The Explorer module integrates benchmark evaluation with downstream biological interpretation. Its benchmark component provides a compact summary of method performance across topology, pseudotime, velocity coherence, computational cost, and prior-information requirements, allowing users to compare methods under different biological and technical settings. The downstream-analysis component offers a richer set of trajectory-based outputs, including Waddington landscapes, driver-gene ranking, gene-density profiles, and gene-trend plots, which jointly support the interpretation of lineage progression and the prioritization of candidate regulators. By connecting quantitative benchmarking with biological exploration in a single interface, Explorer bridges method assessment and hypothesis generation.

### Cafe enables atlas-level cell fate analysis through a divide-and-conquer strategy

Large atlas-scale single-cell datasets are difficult to analyze directly because global fate inference is computationally expensive and often yields noisy, tangled trajectories^[23]^. To address this challenge, Cafe adopt a hierarchical milestone-network framework that implements a divide-and-conquer strategy.

In the gastrulation atlas, coarse- and fine-grained annotations exhibited one-to-many mapping structure (Figure 2a), consistent with StaVia^[24]^. For example, the coarse blood compartment mapped to multiple fine-grained states, including Erythroid1/2/3, Blood progenitors1/2, and their corresponding progenitor cells. A coarse milestone network was first constructed from prior biological knowledge using the projection wrapper to define node- and edge-level subproblems (Figure 2b; divide step)^[25, 26]^. Each subproblem was then solved with a task-specific trajectory method, exemplified by VeloAE for blood cells (Figure 2c; conquer). The local solutions were subsequently merged back onto the coarse scaffold to reconstruct a fine-grained global network (Figure 2d; merge) with the blood section omitted from the main panel because of space constraints; the complete intermediate process is shown in Extended Data Figure 1.

**Figure 2.**
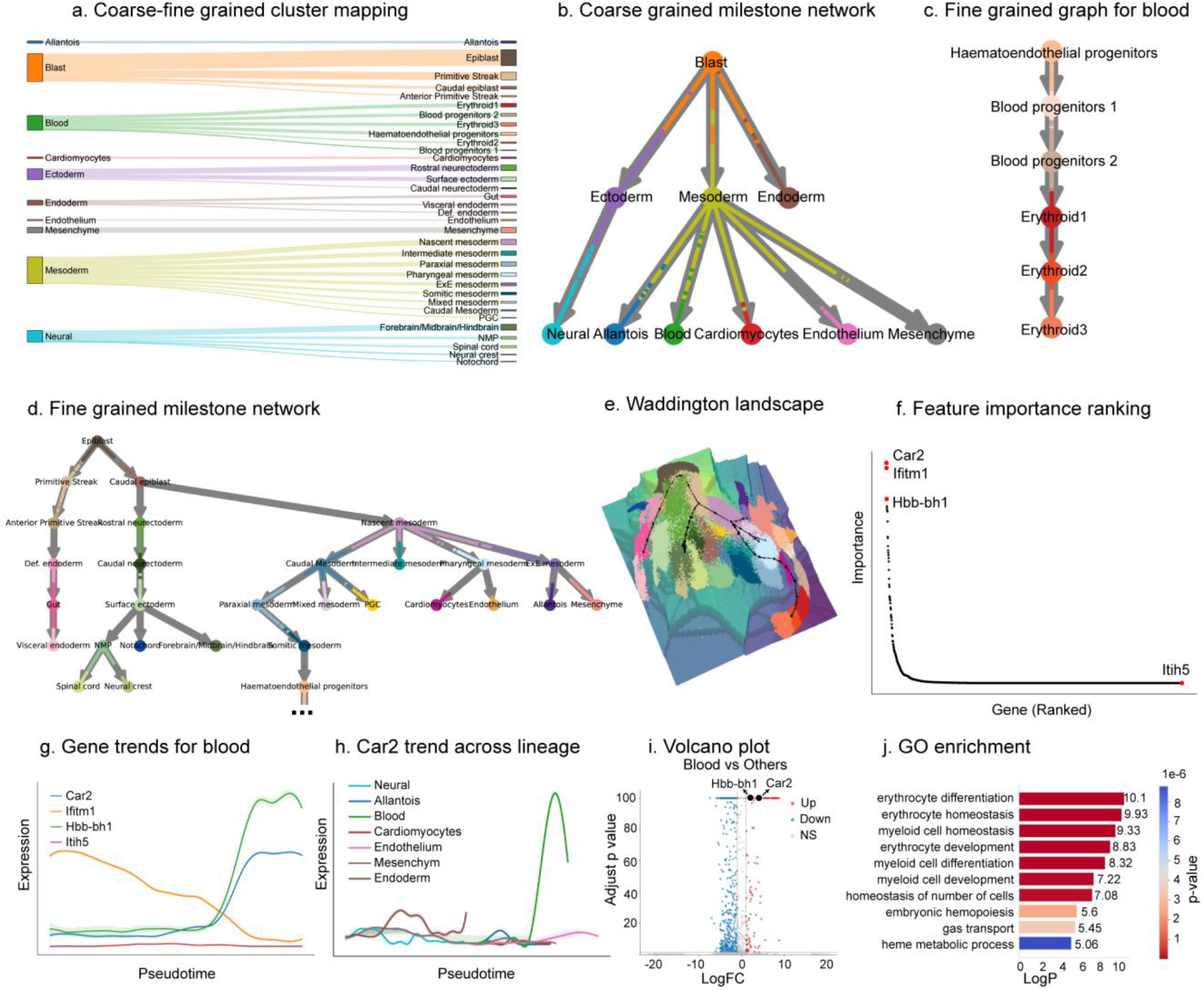
Hierarchical milestone-network analysis enables coarse-to-fine fate inference and driver-gene discovery in a gastrulation atlas. **a**, Mapping between coarse and fine cell annotations (Sankey diagram). **b**, Coarse milestone network used to define subproblems. **c**, Local trajectory inference on split subdatasets, exemplified by VeloAE for the blood compartment. **d**, Reconstructed fine-grained milestone network after merging local solutions. **e**, with x and y denoting UMAP coordinates and z denoting smoothed pseudotime. **f**, Feature-importance ranking of candidate driver genes. **g**, Pseudotime trends of candidate genes in the local blood trajectory. **h**, Global lineage-resolved trends validating lineage specificity. **i**, Volcano plot highlighting candidate genes in differential expression analysis. **j**, GO enrichment of candidate-associated programs by MetaScape^[29]^, with significant erythropoiesis/hematopoiesis terms.

The resulting topology yielded a coherent waddington landscape (Figure 2e)^[27]^, where x and y corresponding to UMAP coordinates and z is the smoothed pseudotime. The same structure further supported driver-gene prioritization: feature-importance ranking identified Car2, Ifitm1, and Hbb-bh1 as top candidate drivers^[28]^, whereas Itih5 served as a control (Figure 2f). In the local blood trajectory, candidate genes tracked fate progression, whereas the control gene showed no clear association with pseudotime (Figure 2g). At the global lineage level, Car2 remained specifically associated with the blood branch (Figure 2h), and differential-expression analysis further supported its lineage specificity (Figure 2i). GO enrichment of the top 25 features highlighted erythropoiesis- and hematopoiesis-related pathways (Figure 2j), supporting Car2 as a putative driver of blood-lineage progression in the gastrulation atlas.

To translate this coarse-to-fine logic into a reproducible workflow, the corresponding divide-and-conquer procedure is formalized in the **Algorithm 1** and **Algorithm 2**.

### Cafe benchmark trajectory among various methods

To benchmark Cafe across heterogeneous trajectory inference results, representative methods spanning velocity-based, linear, cluster-based, probability-based, directed, graph-based, and lineage-based approaches were standardized into a common milestone-network representation. This unified interface enabled direct comparison of topology, pseudo-time correlation, velocity coherence, resource usage, and dependence on prior information within the same evaluation framework. The corresponding per-method trajectory reconstructions are shown in Extended Data Figure 3, and the integrated benchmark summary is reported in Figure 3. Overall score is calculated by several efficient metrics, illustrated in method chapter.

**Figure 3.**
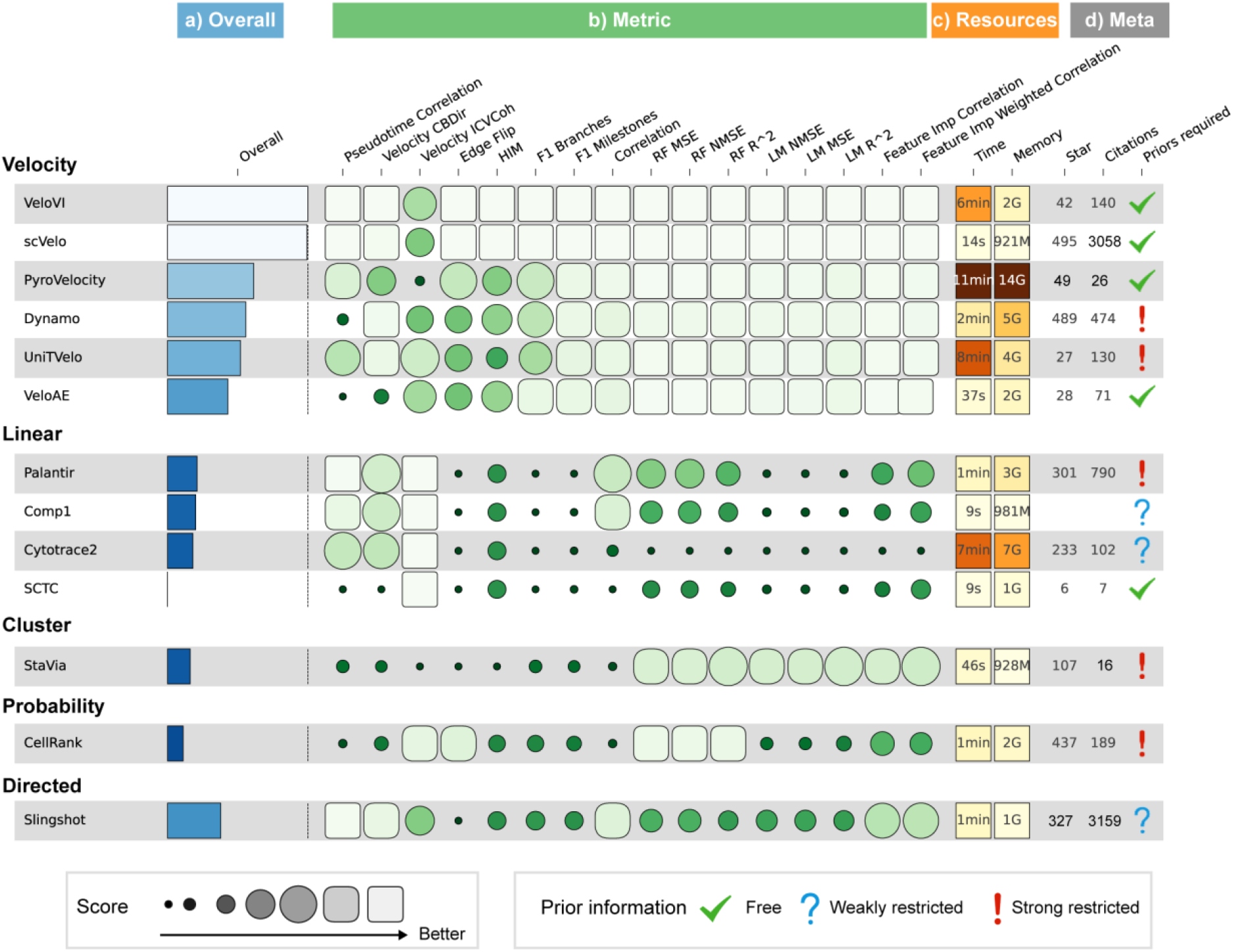
Cafe benchmark trajectory among various methods in the pancreas dataset, visualized by funkyheatmap^[31]^. Rows correspond to methods and are grouped by trajectory type. **a**, Overall score summary. **b**, Normalized performance across topology, pseudotime, velocity coherence, branch assignment, position prediction, feature importance, runtime, and meta-information categories. **c**, Runtime and memory usage. **b**, Meta-information, including GitHub star count, citation count, and prior-information requirements. Circle size and color intensity encode normalized performance, with larger and darker circles indicating better scores. Runtime and memory values are shown in the resource panel; zero values indicate unavailable or failed measurements.

**Figure 4.**
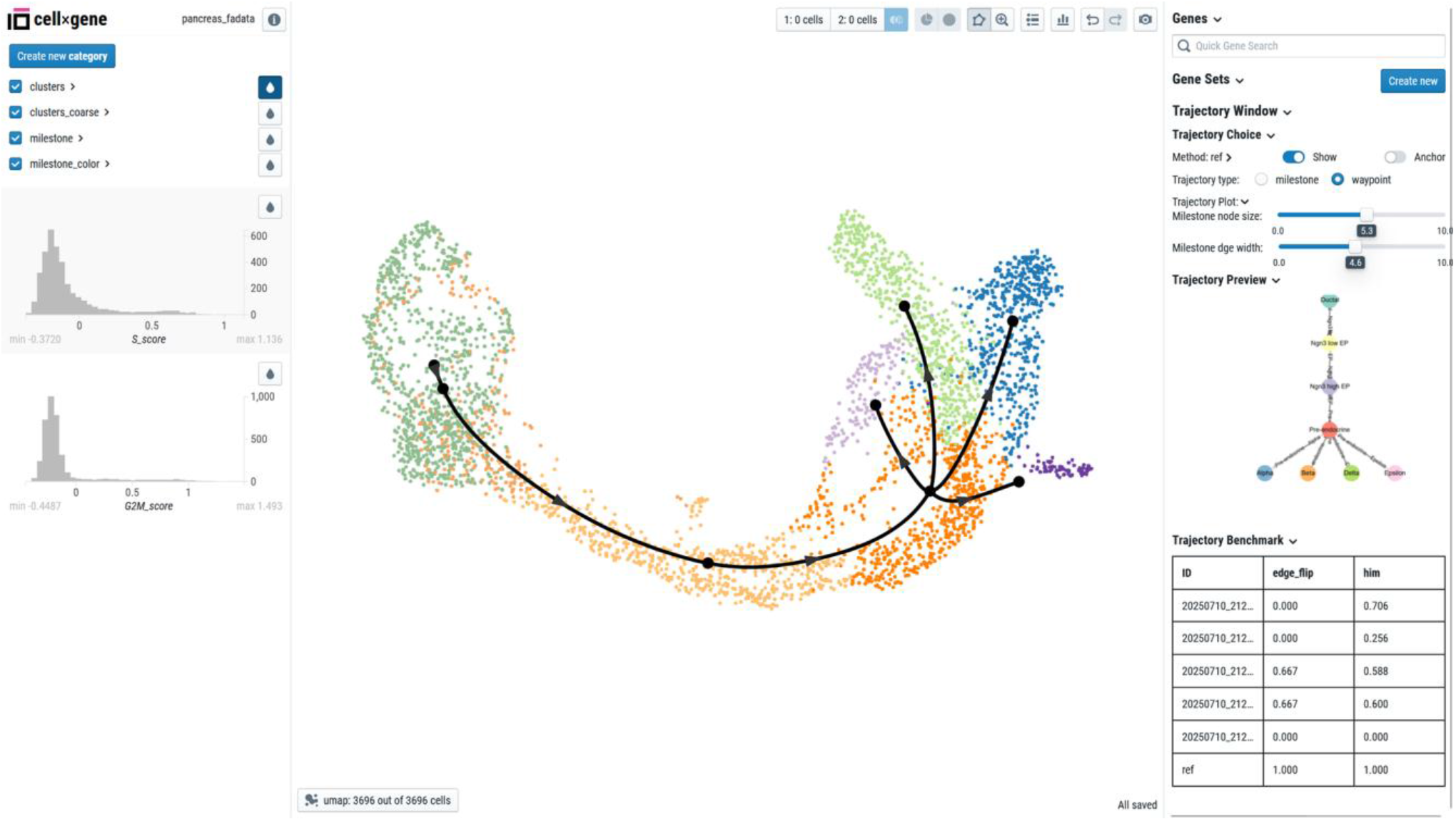
Cellxgene-Cafe enables interactive visualization and evaluation of cell fate trajectories. The left panel displays annotation layers and score distributions for rapid dataset inspection. The center panel overlays the inferred lineage topology onto the single-cell embedding, enabling direct assessment of whether the recovered trajectory follows the underlying cellular manifold. The right panel provides gene and gene-set search, trajectory-type selection, rendering controls for milestone and waypoint views, a live trajectory preview, and a benchmark table.

Based on the pancreas dataset across various methods, the benchmark revealed clear trade-offs rather than a single universally optimal method. scVelo reconstructed a smooth flow field that followed the major developmental continuum and preserved the branching geometry of the reference manifold, consistent with strong velocity-coherence and topology-related scores. By contrast, SCTC recovered an inverted linear backbone, whereas StaVia produced a more complex but redundant milestone network. Both deviated from the expected biological progression and therefore received lower scores. Slingshot recovered a compact and nearly linear backbone, which is well suited to simple trajectories but tends to un-derrepresent branching complexity in atlas-scale differentiation, leading to intermediate branch-recovery and topology-consistency scores^[30]^. More generally, methods that better preserved branch continuity and local geometry achieved higher scores in topology, pseudotime, and velocity-related metrics, whereas simpler linear or directed methods were computationally efficient but less expressive for branching structures. Together, these results indicate that method choice should be matched to the biological structure of the dataset and to the amount of prior information available. The benchmark can be readily extended to additional datasets wrapped in the FateAnnData data structure.

To complement the benchmark, Meta information such as descriptors, citation count and GitHub star count were reported, which capture scholarly visibility and software-community adoption and as of April 17, 2026, respectively. These measures were kept separate from performance scores because they reflect usage and maintenance rather than predictive accuracy. Prior information requirements are categorized as free, weakly restricted, or strongly restricted. Free methods require no prior information; weakly restricted methods can optionally use inputs such as cluster labels or a start cell; strongly restricted methods require prior information to run. This classification helps distinguish methods by deployment flexibility and biological guidance.

### Cafe is equipped with an interactive analysis platform

To bridge the gap between computational trajectory inference and biological interpretation, we developed Cellxgene-Cafe, a browser-based interactive analysis platform built on the scalable cellxgene architecture and seamlessly integrated into Cafe^[18]^. Rather than requiring users to alternate between command-line execution, static figures, and separate downstream tools, Cellxgene-Cafe consolidates trajectory inspection, benchmark evaluation, and biological interpretation into a single web environment. This design lowers the barrier to cell fate analysis by enabling users without programming expertise to complete trajectory exploration, assess quality, and interpret results directly in the browser.

The platform overlays the inferred lineage topology on the single-cell embedding, allowing users to visually assess whether the recovered trajectory follows the underlying cellular manifold and whether key branching events are biologically plausible. The central panel provides an immediate view of the predicted lineage on the embedding, while the side panels support dataset inspection, gene and gene-set search, trajectory selection, and interactive adjustment of rendering parameters such as milestone size and edge width. Importantly, the platform couples visualization with real-time benchmarking, so that topological fidelity can be evaluated alongside the displayed trajectory rather than after separate post-processing. By integrating no-code exploration, quantitative assessment, and expression-based interpretation in one interface, Cellxgene-Cafe enables rapid validation of inferred cell fates and facilitates hypothesis generation from trajectory results.

## Methods and Materials

### Dataset

Cafe provides a unified data layer for both externally curated benchmark datasets and built-in case-study datasets. The framework directly accommodates AnnData obejct and Dynverse-formatted benchmark inputs while preserving reference trajectories, cell annotations, and other trajectory-aware metadata in a common FateAnnData representation. This allows datasets from different sources to be handled consistently while retaining the gold-standard labels required for fair trajectory evaluation.

The dataset collection summarized in Extended Data Figure 3 spans representative developmental programs, including pancreas, CellRank-based pancreas, erythroid lineage, bone marrow, and hematopoiesis. These datasets cover linear, bifurcating, and multi-branch topologies, thereby providing diverse benchmark settings for evaluating trajectory inference methods under different biological and technical conditions. In each case, the reference trajectory serves as the gold-standard label, and the corresponding single-cell embedding provides the geometric context for assessing whether reconstructed lineages recover the expected developmental structure.

### Wrapper description and visualization

To standardize heterogeneous trajectory outputs, as shown in Extended Data Figure 4a, Cafe implements a milestone wrapper abstraction that represents each trajectory as a directed milestone network *T* = (*G, P*), where *G* = (*V, E*) denotes the milestone network graph, *V* s the set of discrete milestones, *E* ⊆ *V* ×*V* is a weighted directed graph, and *P* stores cell progression along trajectory edges. *P*_*c*_ is the projection of cell *c* on its corresponding edge (*u, v*) .In this representation, milestones encode discrete cell states, edges capture lineage transitions, and divergence regions describe local branching events. Waypoints are sampled at regular intervals along milestone edges to discretize the trajectory manifold and provide anchors for visualization and metric computation. This abstraction decouples trajectory topology from method-specific output formats and provides a common coordinate system for downstream analysis.

#### Directed wrapper

The milestone network and cell percentage provide an abstract representation of cell trajectories as shown in Extended Data Figure 4a, Milestones represent key nodes in the trajectory, forming a graph structure, whereas percentages indicate the strength of cell assignments to trajectory edges. Waypoints are sampled at equal intervals along milestone edges, facilitating metric computation and mapping between topological structures and cell embeddings.

As illustrated in Extended Data Figure 4b, geodesic distance is computed on the milestone network rather than in euclidean space. The geodesic distance based on milestone network between cell *a* and *b* is defined:

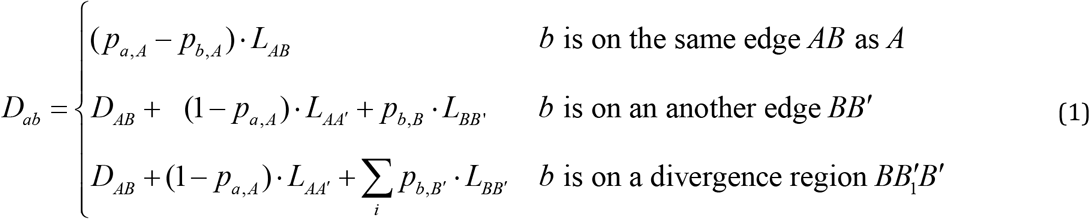

where*p*_*a,A*_ is the progression value and *L*_*AB*_ is the length of edge (*A, B*) . For points on different edges, the distance is obtained by adding the residual distances to their nearest milestones and the shortest path between those milestones; when one point lies in a divergence region, the distance is computed by weighting the outgoing branches according to its progression values. In this way, the directed wrapper preserves lineage orientation while remaining compatible with both tree-like and graph-like topologies.°

To map the abstract topology back into the embedding space, Cafe computes a cell-to-waypoint distance matrix. *D* ∈[0, +∞]^*N*×*K*^, where *N* and *K* are the numbers of cells and waypoints, respectively. The contribution of cell *i* to way-point *j* is normalized as

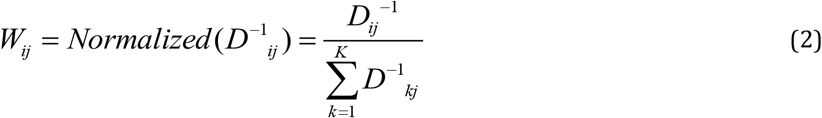

The coordinate of waypoint *j* is defined as the weighted centroid

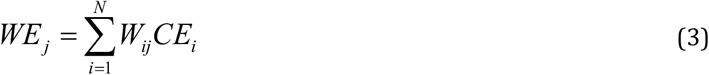

*CE*_*i*_ denotes the low-dimensional embedding coordinate of cell *i* . The weighted projection ensures that the displayed trajectory faithfully follows the underlying data manifold while preserving the topology defined by the milestone network. The remaining seven wrappers in Figure 1b can be transformed into the direct wrapper shown in Extended Data Figure 4a through corresponding wrapper methods. These wrappers convert diverse data structures to standardized representation consisting of milestone network, cell percentage, progression values and divergence region(optional). The details of these wrapper methods shown below.

#### Linear wrapper

The linear wrapper takes a one-dimensional pseudotime sequence as input and constructs a single-edge milestone network from start to end, where each cell’s progression percentage corresponds to its normalized pseudotime value.

#### Cycle wrapper

The ycle Wrapper likewise operates on pseudotime but constructs a cyclic topology with three milestones (A→B→C→A), uniformly distributing the pseudotime across the three edges.

#### Probability wrapper

Probability Wrapper receives an end-state probability matrix and an optional pseudotime sequence as input. When only a single terminal state exists, it degenerates to a linear trajectory. For multiple terminal states, it constructs a star-shaped milestone network centered on a virtual starting point, with each terminal state corresponding to a radial edge. Each cell’s progression percentage along each edge is determined by its terminal-state probability with pseudotime, thereby capturing its differentiation propensity toward distinct fates within the divergence region.

#### Cluster wrapper

The cluster wrapper maps cells to milestone edges using an inter-cluster connectivity graph together with cell cluster labels. Rather than requiring a continuous embedding or cell-level transition vector, it treats cluster relationships as the basic units of topology and positions each cell on the edge associated with its cluster. This design is particularly useful for methods that naturally produce cluster-based outputs or for datasets in which coarse annotations are more reliable than fine-grained progression estimates. By converting discrete cluster structure into a milestone network, the wrapper preserves the major lineage scaffold while remaining compatible with the unified progression representation used throughout Cafe.

#### Projection wrapper

The projection wrapper computes cell positions by geometric projection onto milestone edges using the embedding coordinates of both milestones and cells. For each cell, all edges are examined, and the edge with the minimum projection distance is selected as the assignment. The projection ratio is calculated from the vector dot product, out-of-bound projections are clamped to the edge boundaries, and the projected point together with its distance to the cell is then determined sequentially. The resulting edge assignment and progression value provide a smooth mapping from low-dimensional embeddings to the milestone network.

The detailed geometric projection algorithm operates as follows, as illustrated in Extended Data Figure 4d. For a given cell *p*, the algorithm iterates over all edges in the milestone network and selects the edge with the minimum projection distance. Taking edge *XY* as an example:

1. Compute progression percentage: The projection position ratio is calculated using the vector dot product as following:

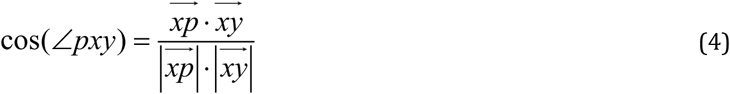

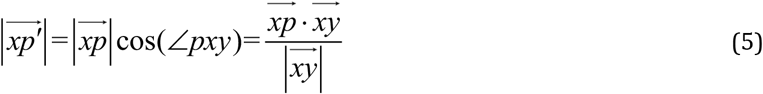

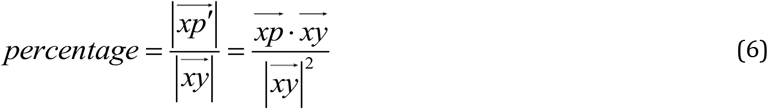
2. Boundary clamping: If the projection falls outside the edge boundaries (as shown for *p*_2_ and *p*_3_ in the figure), the percentage is clamped to the interval [0,1]:

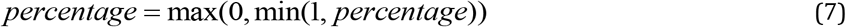
3. Compute projection coordinates and Distance: Determine the projection point *p*^′^ and calculate the projection distance *dist* .

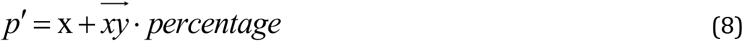

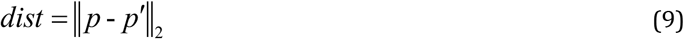
4. Select nearest edge: After iterating through all edges, the edge with the minimum projection distance is assigned to the cell.

Because this geometric operation is independent of the original inference method, it can also be reused in the velocity and lineage wrappers, as well as in manual trajectory construction.

#### Graph wrapper

The graph wrapper starts from a cell-level connectivity graph, identifies key cells to retain as milestone candidates, and then extracts a simplified backbone network before remapping all cells onto the reduced trajectory. To control the degree of simplification, Cafe implements a two-stage strategy with a tunable pruning threshold. In branch pruning, the threshold is multiplied by the graph diameter to obtain an absolute cutoff, and terminal branches shorter than this cutoff are collapsed into the nearest backbone node. In chain simplification, relay nodes with degree two in undirected graphs or with in-degree and out-degree both equal to one in directed graphs are removed, and adjacent edges are merged with accumulated weights. Increasing the pruning threshold therefore produces a sparser network with fewer milestone nodes, allowing the user to trade structural detail for topological robustness.

The graph wrapper takes a cell-level connectivity graph, identifies key cells to retain as milestone candidates, extracts a backbone network through graph simplification, and then maps all cells onto the simplified milestone network. For the graph simplification process, we implemented an optimized two-phase strategy with tunable parameters:

1. Branch Pruning: A pruning threshold (ranging from 0 to 1) is specified, which is multiplied by the graph diameter to obtain the absolute threshold. The algorithm iteratively searches inward from leaf nodes; if a branch’s total length falls below the threshold before reaching a bifurcation point, the nodes along that branch are collapsed onto the nearest backbone node, and the cells residing on these edges are remapped accordingly.
2. Chain Simplification: Nodes with degree 2 in undirected graphs, or nodes with both in-degree and out-degree equal to 1 in directed graphs, are classified as relay nodes and subsequently removed. Adjacent edges are merged with accumulated weights, and cells originally positioned on the removed edges are proportionally remapped onto the newly merged edge based on their relative distances.

As illustrated in Extended Data Figure 4e, Increasing the pruning threshold therefore yields a sparser network with fewer milestone nodes, allowing the user to trade structural detail for topological robustness.

#### Velocity wrapper

The velocity wrapper is designed to convert the outputs of RNA velocity methods into a standardized milestone network representation. Cafe first applies PAGA to the velocity graph, and the resulting transition confidence matrix is used to define directed inter-cluster connectivity. Each cell cluster is treated as a milestone node, with node positions defined by the centroid of all cells in the cluster in the low-dimensional embedding. For simplicity, all milestone edges are assigned unit length, after which individual cells are mapped onto the constructed network through the same geometric projection procedure described above. This allows velocity-informed fate dynamics to be represented within the same topological framework as pseudotime-, graph-, and probability-based trajectories.

#### Lineage wrapper

The Lineage wrapper takes a probability matrix as input, illustrated by Extended Data Figure 5a, where rows represent individual cells and columns correspond to terminal states. This input format is identical to that of the probability wrapper described above. However, the star-shaped network topology generated by the probability wrapper is overly simplistic and fails to recover the complete tree-like lineage structure, as illustrated in Extended Data Figure 5b. To address this limitation, we developed two novel strategies for the lineage wrapper: graph fusion and hierarchical clustering.

The core algorithm first computes a cluster-level probability matrix by averaging cell probabilities within each cluster (Extended Data Figure 5c). The cluster with the smallest summed probability across all terminal states is designated as the root milestone. Based on this cluster probability matrix, a milestone network is then constructed using either of the two proposed strategies described below. Cells are subsequently projected onto the edges in the probability space to obtain milestone progressions. Additionally, each branching node in the network defines a divergence region.

Graph Fusion Strategy: The first strategy employs a graph-based path fusion approach (Extended Data Figure 5d). For each terminal state, a lineage string is constructed by first excluding clusters associated with other terminal states and then sorting the remaining clusters in ascending order of their probability values for that specific terminal state. These lineage strings are then converted into a weighted consensus graph by sliding a window of size two across each string, such that each consecutive cluster pair forms a directed edge. Edge weights are incremented by one for each occurrence across all lineage strings. The final milestone network is obtained through graph pruning, which eliminates cycles and retains only the highest-weighted edges using a breadth-first traversal from the root.

Hierarchical Clustering Strategy: The second strategy leverages hierarchical clustering based on the probability matrix features Extended Data Figure 5e). Terminal states are iteratively merged according to their similarity in the probability space until a single branch point remains, which is then connected to the root. The resulting dendrogram is inverted to yield a directed milestone network emanating from the root toward the terminal states.

Extended Data Figure 5f demonstrates the application of the star, graph fusion, and hierarchical clustering strategies on the CellRank result for pancreas dataset. Both proposed strategies produce milestone networks that more closely approximate the expected reference topology compared to the star-shaped network. Notably, each strategy exhibits distinct characteristics: graph fusion may occasionally omit intermediate clusters during the pruning process, whereas hierarchical clustering invariably produces strictly binary tree structures.

#### Wrapper visualization

Representative wrapper outputs are summarized in Extended Data Figure 6, where both the original method-specific views and the standardized milestone-network views are shown side by side. These examples illustrate that Cafe preserves the topological structure of diverse trajectory inference methods while providing a consistent visual language for comparison, interpretation, and downstream analysis.

### Cell fate prediction methods

Cafe provides a unified interface for cell fate prediction methods, allowing heterogeneous trajectory inference outputs to be converted into a common milestone-network representation. The current implementation covers representative methods across multiple trajectory types:

- Directed: PAGA^[32]^
- Linear: **Comp1**, SCTC, Cytotrace2^[12]^, Palantir^[11]^.
- Cycle: **Angle**.
- Cluster: **ClusterMST**, StaVia^[24].^
- Projection: **ProjectionMST**.
- Graph: **GraphMST**.
- Velocity: CellDancer^[33]^, CellRank^[34]^, Dynamo^[14]^, PyroVelocity, scVelo^[13]^, UniTVelo^[35]^, VeloAE^[36]^, VeloVI^[37]^.
- Lineage/Probability: CellRank^[34]^, Palantir^[11]^, **StateComp**.

In the above list, the bolded methods are baseline for corresponding wrapper. Methods that share the same output type, such as CellRank, can be routed to multiple wrappers depending on the available input and intended analysis mode. In addition to Cafe-native wrappers, the framework also supports direct invocation of Dynverse-compatible methods, enabling established methods such as Slingshot and other Dynverse implementations to be benchmarked within the same evaluation pipeline.

Rather than treating these methods as isolated implementations, Cafe standardizes them through wrapper-specific input and output conventions. This design makes it possible to compare methods with different assumptions, data requirements, and topological expressiveness under a common analysis framework. In the benchmark presented in this study, only representative methods are shown for each wrapper category, but the framework is extensible and can incorporate additional methods by updating the wrapper mapping and backend configuration.

### Workflow Optimization

Cafe reorganizes the conventional workflow of cell fate analysis around a unified milestone-network model and its associated operations, thereby bridging upstream data preparation, core trajectory inference, and downstream biological interpretation in a single framework. Compared with the standard workflow, the Cafe pipeline reduces redundant data conversion and environment switching while improving compatibility with the Scverse and Scanpy ecosystems. In the upstream stage, Cafe first evaluates candidate embeddings for trajectory analysis using structure-aware criteria, such as Cluster Silhouette and Strip Score, to identify embeddings that simultaneously preserve cell-type separation and developmental continuity. When reference annotations or prior lineage information are available, users can further guide trajectory construction through projection-based annotation; otherwise, label-free trajectory inference methods can directly provide the necessary topological structure. In the core stage, heterogeneous trajectory inference methods are standardized into a common milestone-network representation through the wrapper system, enabling consistent comparison across methods with different output formats and biological assumptions. In the downstream stage, the unified milestone network is decomposed into lineages for further analysis, including differential-expression testing and lineage alignment, which facilitates driver-gene discovery and biological interpretation. The complete optimized workflow is summarized in Extended Data Figure 7.

### Backend

Cafe supports four interchangeable execution backends: python function, conda, cafe docker, and Dynverse docker. The python function backend runs native methods directly in the cafe environment, the conda backend isolates methods with dedicated dependencies, the cafe docker backend packages native implementations into reproducible containers, and the Dynverse docker backend reuses curated Dynverse implementations without reimplementation. By abstracting execution through a single interface, Cafe enables heterogeneous trajectory methods to be run consistently while reducing dependency management overhead and preserving extensibility.

### Evaluation metric description

Cafe evaluates trajectory inference from complementary perspectives, covering embedding quality, temporal ordering, velocity coherence, topology preservation, cluster agreement, pairwise distance consistency, state-assignment accuracy, and feature relevance. The metrics introduced in this work are summarized in Extended Data Figure 8, while the remaining metrics follow Dynverse-style benchmarking. Together, these measures provide a balanced assessment of geometric fidelity, topological correctness, and biological interpretability.

#### Embedding Metric

The embedding metric evaluate both the upstream embedding, before trajectory inference, and the core trajectory result, after inference. For the pre-trajectory stage, the selected embedding should separate cell populations and preserve a continuous, lineage-like geometry. We therefore use cluster silhouette to quantify cluster separability and strip score to assess whether the embedding forms a strip-like manifold. Strip score is computed from the minimum spanning tree of the embedding neighborhood graph and increases when the embedding better supports trajectory reconstruction.

For pre-trajectory embedding evaluation, the ideal embedding should satisfy two criteria: it should exhibit clear clustering of distinct cell types and form a continuous, strip-like structure representing the developmental lineage. Cluster silhouette use embedding as features and cluster as label to calculate silhouette coefficient, which is classical metric for unsupervised cluster task. Strip score assesses whether the embedding forms continuous and strip-like structure. As illustrated by Extended Data Figure 8a, firstly, MST (black line) should be found on the cell graph (grey line), constructed by normalized embedding feature. The diameter of the MST (red line) is then divided by the total number of nodes to obtain the strip score as formula (10). In the worst case, when the embedding distribution is close to spherical, the diameter of the corresponding minimum spanning tree is very small. Among the three embedding methods shown in Extended Data Figure 8b, UMAP has a slightly lower cluster silhouette than PCA but is selected for downstream analysis because it yields a higher strip score and is therefore more effective for trajectory inference. The numerical results are consistent with the visual inspection, demonstrating the usefulness of the metric.

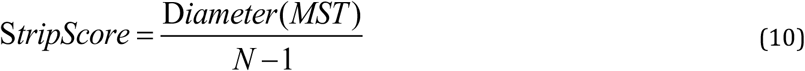

For post-trajectory embedding evaluation, cells that are closer to the starting cell in the embedding space should be in the early stages of differentiation. The distance from the starting cell in embedding space is obtained by calculating the shortest path length on the cell neighbor graph generated by embedding space. The differentiation stage of cells is calculated using pseudo time based on milestone network structure. The metric GDPC (Geodetic distance pseudotime correlation) evaluates the pearson correlation between the pseudo time and distance, two one-dimensional values. In Extended Data Figure 8B, the result of “Ref” trajectory is the most accurate, “trajectory1” generates an incorrect branch at the end, and “trajectory2” generates a reverse trajectory. These three results correspond to the best, medium, and worst cases, respectively. They are also accurately reflected on the visualization and metric value.

#### Pseudotime metric

Pseudotime Correlation (PC) quantifies the agreement between predicted and reference pseudotime by calculating the pearson correlation between the predicted and reference pseudotimes, as shown in Equation(11)

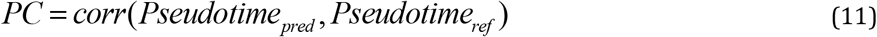

To extend this metric to complex trajectory models (e.g., graph-based or velocity-based trajectories), the trajectory structure transfers to linear pseudotime values through the following steps, as shown in Extended Data Figure 8C:

- Start milestone setting. A root milestone is specified for the milestone network, typically derived from prior information.
- Milestone pseudotime calculation. The shortest geodesic distance from the root to every other milestone is calculated using Dijkstra’s algorithm on the milestone network.
- Cell pseudotime calculation. The pseudotime for each cell is computed as the weighted sum of the associated milestones, based on the cell’s projection onto the trajectory edges.

#### Velocity metric

To quantitatively evaluate the accuracy and robustness of the inferred RNA velocity, we employed two established metrics: Cross-Boundary Direction Correctness (CBDir) and In-Cluster Velocity Coherence (ICVCoh)^[36]^.

CBDir assesses whether the inferred velocity correctly points towards the expected future state during cell differentiation. We focus on the boundary cells, that are expected to transition from cluster *A* to cluster *B*, These boundary cells are defined as the subset of cells in *A* that possess nearest neighbors in cluster *B*, as shown in Eq. (12):

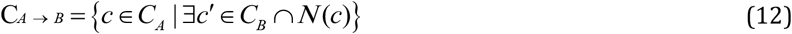

For a given cell *c*, CBDir is calculated as the average cosine similarity between its velocity vector *v*_*c*_ and the displacement vectors *x*_*c*_^′^ − *x*_*c*_ pointing towards its neighbors *c*^′^ in the target cluster *B* . A higher CBDir score indicates that the velocity field correctly guides cells across the boundary into the subsequent differentiation stage. as defined:

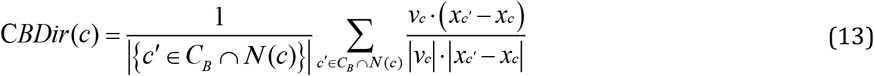

ICVCoh measures the local smoothness and consistency of the velocity field within a specific cell state. It is defined as the average cosine similarity between the velocity *v*_*c*_ of a cell *c* and the velocities *v*_*c*_^′^ of its neighbors within the same cluster. High coherence implies a stable and continuous differentiation flow within the cell population.

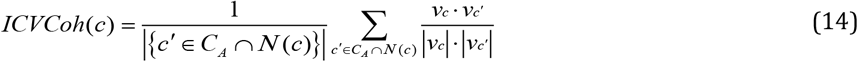

To extend these velocity-based metrics for benchmarking various trajectory inference methods implemented as wrapper in the cafe framework, we developed a strategy to generate pseudo-RNA velocity from the inferred milestone networks. Specifically, within a reduced dimensional space such as UMAP, we assigned velocity vectors to cells located along the edges of the milestone network. For cells residing on a milestone edge, velocities are explicitly set as the unit vector aligned with the direction of that edge. For cells in divergence region, velocities are the weight sum of unit vectors of related edge. This approach allows direct comparison of the topological correctness of trajectory inference methods against standard RNA velocity fields using the CBDir and ICVCoh metrics. Velocity generation and metric case are shown in Extended Data Figure 8D. What’s more, the cluster edges can also be extracted from the ref trajectory milestone network (gold standard label).

#### Topology metric

Topological metrics are employed to assess the structural similarity between the reference and predicted milestone networks, independent of the specific positions of individual cells.

##### Isomorphism

This metric determines whether the reference and predicted network structures are topologically identical. It is directly implemented using the “*networkx*.*is_isomorphic*” function from the *NetworkX* packag^[38]^e, returning 1 if isomorphic and 0 otherwise.

##### Edge Flip

This metric quantifies the topological distance by calculating the minimum number of edge modifications (additions or deletions) required to transform the reference network into a structure isomorphic to the predicted network.

The metric is defined as the number of edge flips divided by the total number of edges in both networks. Because finding the minimum number of flips is equivalent to the **Maximum Common Edge Subgraph** (**MCES**) problem, which is NP-hard, we followed the heuristic algorithm from Dynverse. The algorithm employs an iterative search strategy:

- It starts with the minimum theoretical number of flips, derived from the difference in edge counts between the two networks.
- It incrementally increases the allowed number of flips.
- For each step, it generates all possible combinations of edge additions and removals.
- It applies these changes to the reference network and checks for isomorphism with the predicted network.
- To ensure computational feasibility, the search is constrained by a maximum limit on the number of flips and combinations.

##### HIM (Hamming-Ipsen-Mikhailov)

The HIM metric combines the Hamming distance, which measures local structural differences based on edge matching, and the Ipsen-Mikhailov distance which measures global structural differences based on spectral properties. It provides a comprehensive assessment of topological similarity, normalized to the range [0, 1].

#### Cluster metric

Cluster metrics are employed to assess the structural consistency between the reference trajectory and the predicted trajectory by comparing their induced cell clusters. Milestones and edges are the fundamental building blocks of a trajectory network. Consequently, cells can be intuitively mapped to these entities to form trajectory-specific clusters. As illustrated in Extended Data Figure 8E, in addition to standard reference cluster, cells are grouped to their nearest milestones or highest percentage edges.

This transforms the trajectory evaluation into a supervised clustering comparison task. While standard clustering metrics tools exist^[39]^. We follow dynverse and evaluate the alignment between the predicted clusters *C* and the reference clusters *C*^′^ using **f1 score**. which is the harmonic mean of **recovery** and **revelance**. These components are derived from the maximum Jaccard similarity between the cluster sets^[40, 41]^, defined as follows:

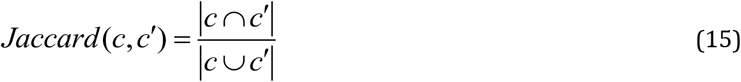

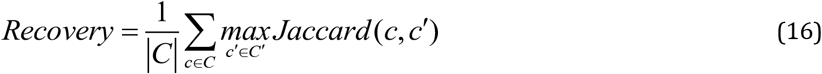

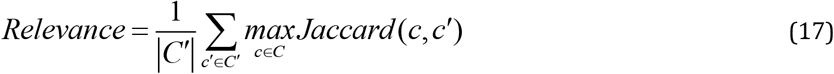

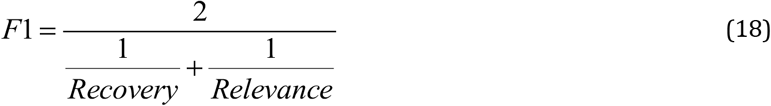

A higher F1 score indicates greater consistency between the predicted trajectory structure and the reference. The metrics corresponding to the two clustering strategies are denoted as *F*1_*milestone*_ and *F*1_*edge*_, respectively.

#### Distance metrics

To evaluate the global structural similarity between the inferred trajectory and the ground truth, we employed a correlation-based metric that compares the geodesic distances between cells and waypoints. This approach is robust to local noise and dimensional scaling differences.

The Correlation Score is defined as the Spearman rank correlation coefficient *ρ* between the flattened vectors of these two distance matrices:

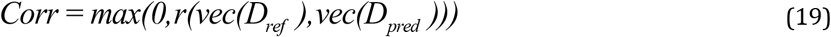

Here, *vec*(.) denotes the flattening operation. We use spearman correlation to assess monotonic relationships, making the metric insensitive to linear scaling or non-linear monotonic transformations of pseudotime units. A higher score, closer to 1, indicates that the predicted trajectory preserves the relative ordering and global connectivity structure of the cells as defined by the reference.

#### Position metrics

Position metrics assess whether the inferred trajectory model correctly maps cells to their biological states compared to a reference trajectory. We treat this as a regression problem: predicting the reference milestone percentages from the inferred milestone percentages.

Let *P*_*ref*_ ∈[0,1]^*N*×*M*^ be the matrix of milestone percentages for *N* cells and *M* milestones in the reference trajectory, and let *P*_*pred*_ ∈[0,1]^*N*×*M*′^ be the corresponding matrix from the inferred trajectory, also referred to as the milestone feature matrix. We construct a multi-label regression model f: *P*_*pred*_ → *P*_*ref*_, in which every milestone corresponds a label. These datasets are split into training (70%) and testing (30%) sets with a fixed random seed for reproducibility. The following regression model are involved:

- Random Forest (RF): A non-linear regressor capable of capturing complex manifold mappings.
- Linear Regression (LM): A linear model to test simple linear relationships.

Mean Squared Error (MSE) evaluate the multi-label regression model performance. Compared with the fixed worst case baseline, in which regression outputs are identical across labels, the less *MSE*, the more *NMSE* (Normalized MSE)illustrated in formula (20). To summarize performance across label, the average *NMSE* ultimately generates *NMSE*_*rf*_

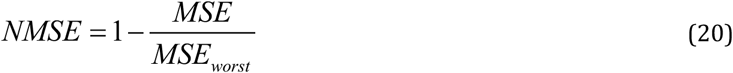

#### Feature metrics

Feature metrics *corr*_*feature*_ assess whether the inferred trajectory recovers biologically meaningful genes associated with progression. We rank genes by their trajectory-related importance scores and compare the ranked profiles between reference and predicted trajectories using correlation- and enrichment-based measures. Correlation evaluates consistency of the overall feature-importance ordering, whereas enrichment tests whether the reference feature set is preferentially concentrated among the top-ranked genes in the predicted trajectory. Together, these metrics quantify the ability of a method to recover candidate driver genes.

#### Overall score

Finally, the overall score integrates the selected metric families into a single benchmark summary, enabling direct comparison across methods and datasets

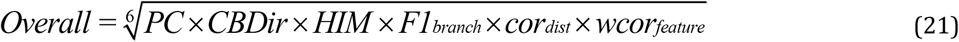

### Divide-and-conquer strategy for atlas-level cell fate prediction

To make the atlas-level cell-fate prediction workflow explicit and reproducible, we formalize the problem definition: Given a global coarse-grained trajectory graph *T*_*gc*_ and a target milestone list *M* = (*m*_1_, *m*_2_,..., *m*_*k*_), the goal is to decompose the dataset into milestone-specific subproblems, apply a cell-fate prediction method *ϕ*_*i*_ to each local subset to obtain a fine-grained local trajectory graph *T*_*lf*_, and then merge the local results back into a global fine-grained trajectory *T*_*gf*_. The divide-and-conquer strategy proceeds in three steps. In the divide step, ells associated with each target milestone *m*_*i*_ are extracted from the full dataset. In the conquer step, the corresponding local subset is processed by *ϕ*_*i*_ to infer the local fine-grained trajectory graph *T*_*lf*_, In the merge step, *T*_*lf*_ is mapped to its corresponding position. Finally, the edge lengths in *T*_*gf*_ are adjusted proportionally to the cell load, or progression density, along each edge, thereby compressing redundant paths and stretching dense developmental transitions.

#### Algorithm 1

Divide-and-Conquer Reconstruction of Atlas-Level Trajectory Graph

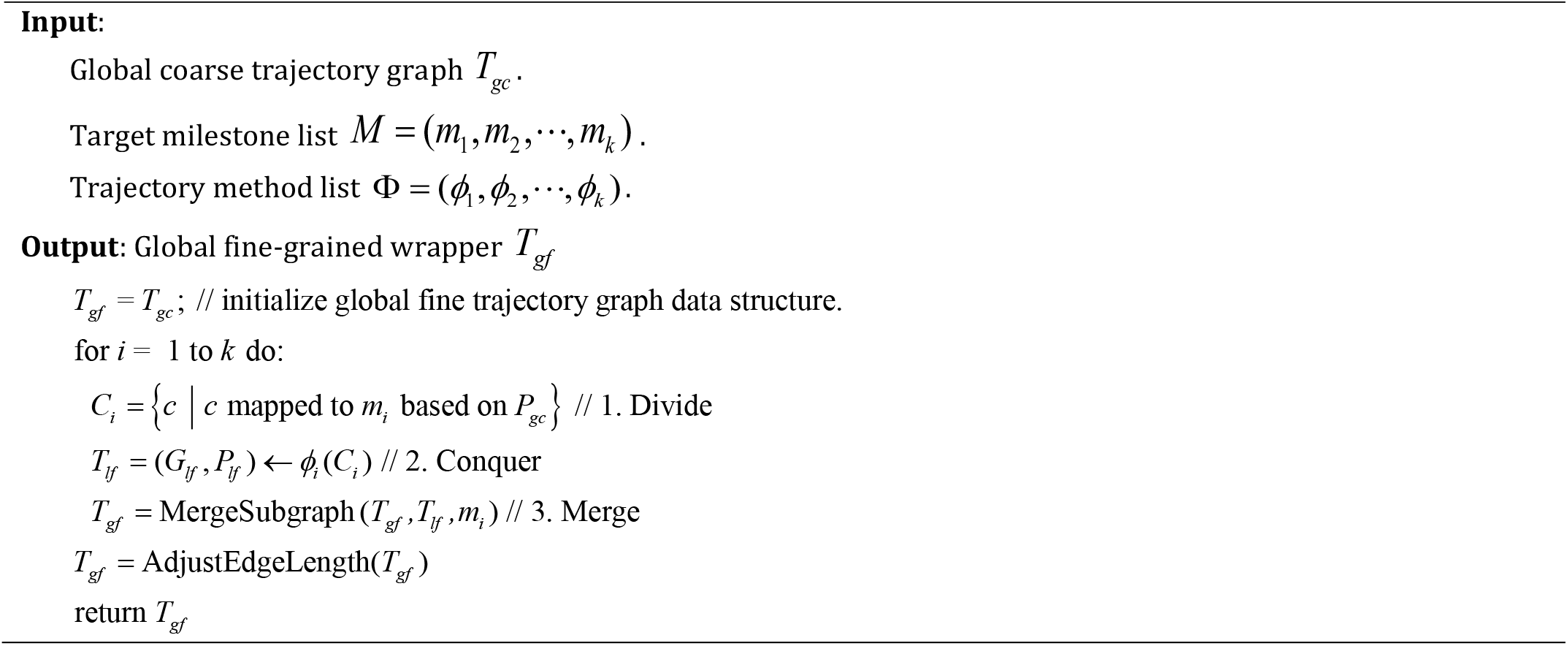

Because of the trajectory graph contains both milestone network *G* and cell progression *P*, the above three key steps require simultaneous adjustment of the two types of data. We therefore describe the subgraph integration procedure separately in Algorithm 2, which is the most difficult sub step. In the algorithm milestone network adjustment need to bridge proper predecessor edges and successor edges, while cell progression should consider how to handle inner, bridged and unrelated cells.

#### Algorithm 2

Subgraph Integration for Global Topology Merging

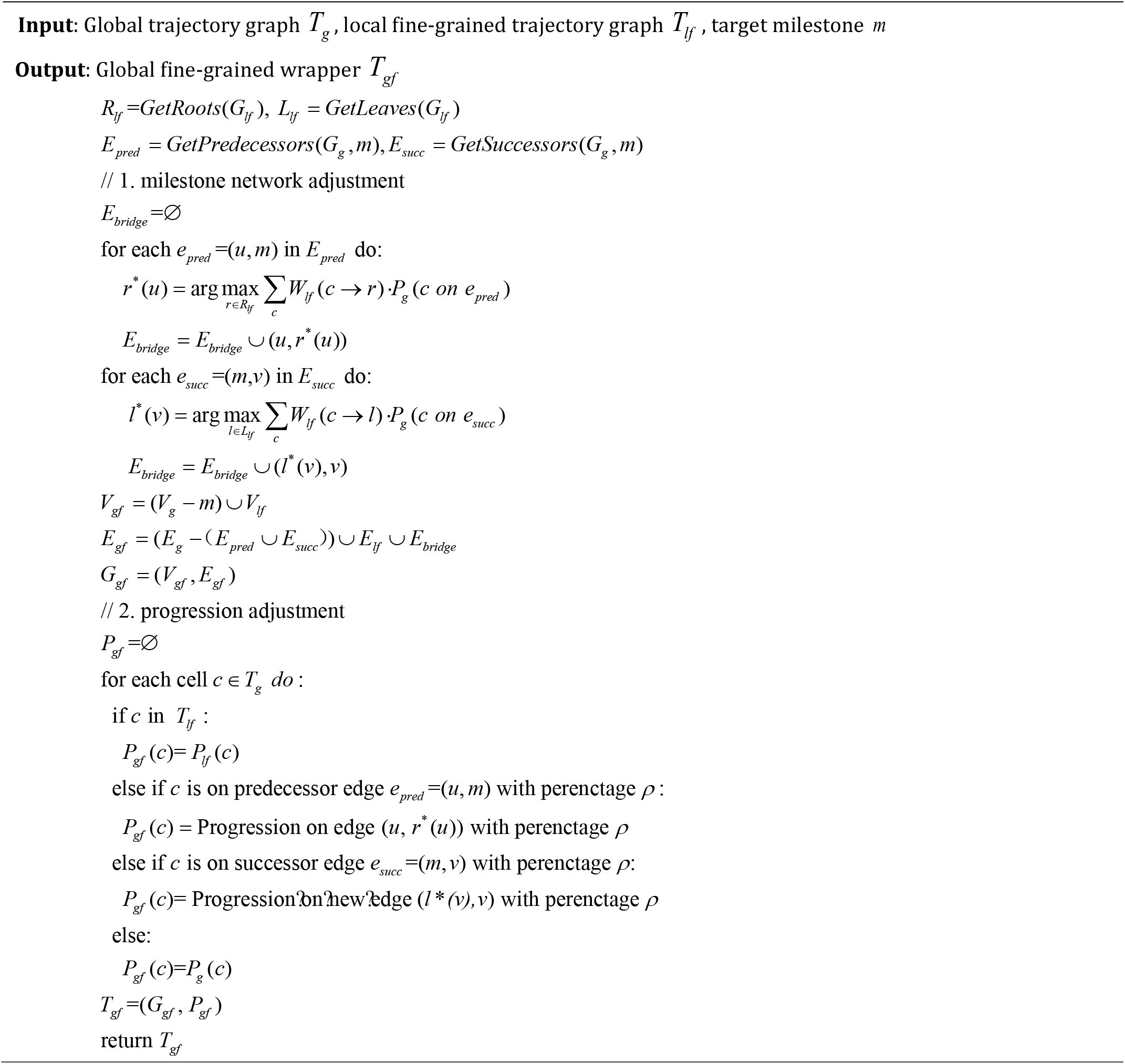

## Disscussion

Cafe is designed as a modular and extensible ecosystem for systematic cell fate analysis, connecting data preparation, trajectory inference, visualization, benchmarking, and biological interpretation within a unified framework. By standardizing heterogeneous outputs through a milestone-network abstraction, Cafe provides a common analytical language for methods comparison while preserving the flexibility required by diverse biological questions and data types. This design makes it possible to move from trajectory reconstruction to interpretation within the same platform, reducing fragmentation across tools and improving reproducibility.

For the data module, future work will focus on expanding support for atlas-scale datasets and additional modalities, including multi-omics^[42]^, spatial transcriptomics and proteomics^[43, 44]^, while also enriching the collection of reference trajectories and gold-standard annotations. Such expansion will make Cafe more representative of the growing diversity of single-cell studies and will strengthen its value as a benchmark-ready analysis platform.

For the method module, we plan to incorporate additional wrappers and broader trajectory representations, including transition-matrix-based formulations that can better capture uncertainty and state-to-state dynamics^[45, 46]^. This will further improve compatibility with emerging trajectory inference methods and allow Cafe to keep pace with the rapidly evolving landscape of cell-fate modeling.

For the explorer module, future development will emphasize three directions. Firstly, Cafe should be extended from basic trajectory-result evaluation to systematic benchmarking linked to open-problem formulations and large-scale cross-dataset comparisons^[47]^. Secondly, downstream analysis will be broadened to include additional interpretation tools, such as driver-gene identification (e.g., BEAM^[48]^ and Gene2Gene^[49]^) and gene regulatory network inference (e.g., SCENIC^[50]^ and CellOracle^[51]^), so that inferred trajectories can be translated into mechanistic biological hypotheses. Thirdly, we envision integrating an AI assistant or agent layer, inspired by systems such as OmicVerse and OmicClaw^[21]^, to help users navigate workflows, recommend methods, and summarize results interactively. Together, these extensions will further lower the barrier to trajectory analysis and reinforce Cafe as a practical platform for cell fate exploration.

In summary, Cafe provides a flexible foundation for trajectory inference and interpretation, with substantial room for future expansion in data coverage, method diversity, and interactive analysis.

## Availability

Cafe source codes are available at https://github.com/HuangDDU/cafe. Tools Cellxgene-Cafe source codes are available at https://github.com/HuangDDU/cellxgene. Document can be accessed by https://cafe-release.readthedocs.io. Pipy package can be accessed by https://pypi.org/project/cafe-release/.

## Acknowledgements

Thanks to all those who maintain excellent databases and to all experimentalists who enabled this work by making their data publicly available.

## Funding

This work has been supported by the National Natural Science Foundation of China (grant nos. 62472344 and 62272065), Natural Science Basic Research Program of Shaanxi Province (No:2025JC-JCQN-094).

## Contributions

All authors contributed to the article. ZH and LY initiated and conceptualized the study. ZH was responsible for developing the wrapper framework and method modules. ZH and HM implemented the metrics and conducted the benchmark experiments. ZH and YP was in charge of the web server development. ZH and CZ conceived and executed the case study. ZH and LY led the writing of the manuscript, which was subsequently reviewed, edited, and approved by all authors. All authors read and approved the final manuscript.

## Competing interests

The authors declare no competing interests.

## Extended data figure

**Extended data figure 1.**
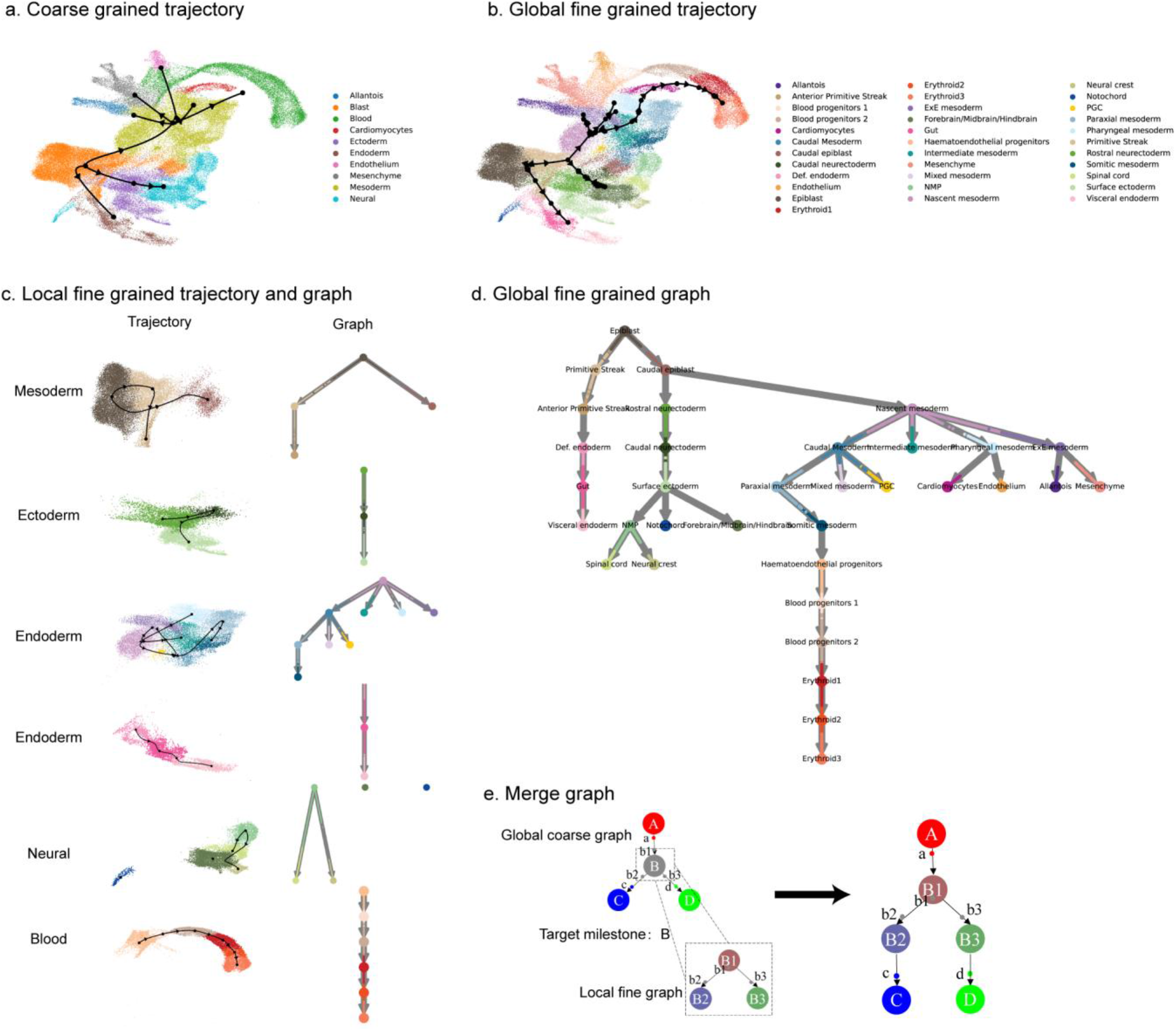
Complete intermediate process and result for gastrulation. **a**, Global coarse-grained trajectory. **b**, Reconstructed global fine-grained trajectory after merging local results. **c**, Multiple local fine-grained trajectories and their corresponding graphs. **d**, Merge graph for fine-grained cluster. **e**, Schematic illustration of the merge operation on the global coarse graph, in which the target milestone B is replaced by the corresponding local fine-grained subgraph to obtain the final global fine-grained graph. Small black dots denote cells, big colored points denote milestones, and arrows indicate the direction of progression.

**Extended data figure 2.**
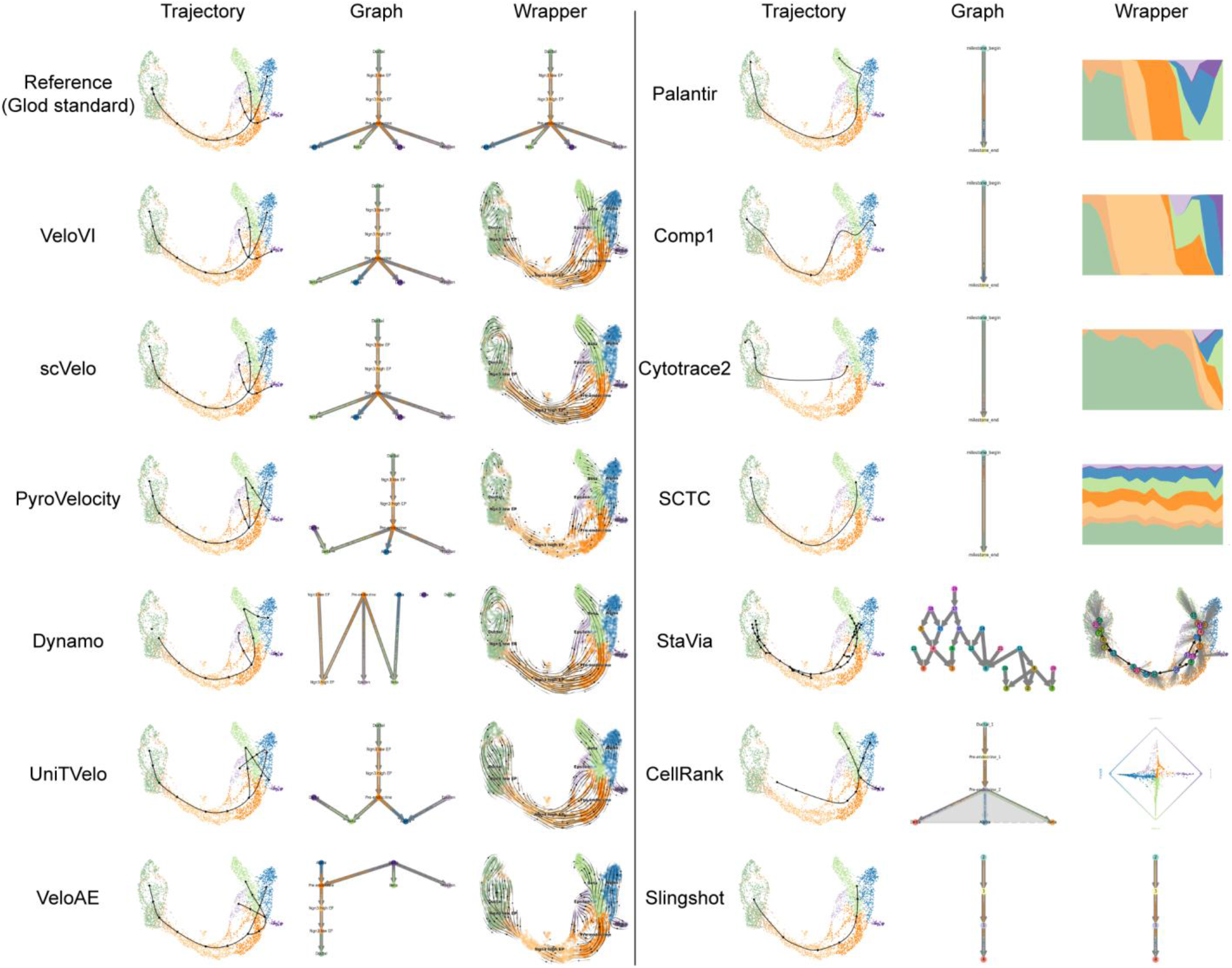
Trajectory-level comparison of benchmarked methods on the pancreas dataset. Each row corresponds to the reference trajectory or one benchmarked method. The left panel shows the inferred trajectory in the shared embedding, the middle panel shows the graph abstraction or milestone graph returned by Cafe, and the right panel shows the wrapper-specific visualization produced by the corresponding backend. Methods are grouped by wrapper type and ordered according to the benchmark summary, highlighting that Cafe preserves a consistent visual language across heterogeneous backends while still allowing method-specific outputs. Representative methods include velocity-based approaches (scVelo, Dynamo, VeloVI, PyroVelocity, VeloAE, UniTVelo), linear or pseudotime methods (Comp1, Palantir, Cytotrace2, SCTC), cluster-based methods (StaVia), probability-based methods (CellRank), and directed methods (Slingshot).

**Extended data figure 3.**
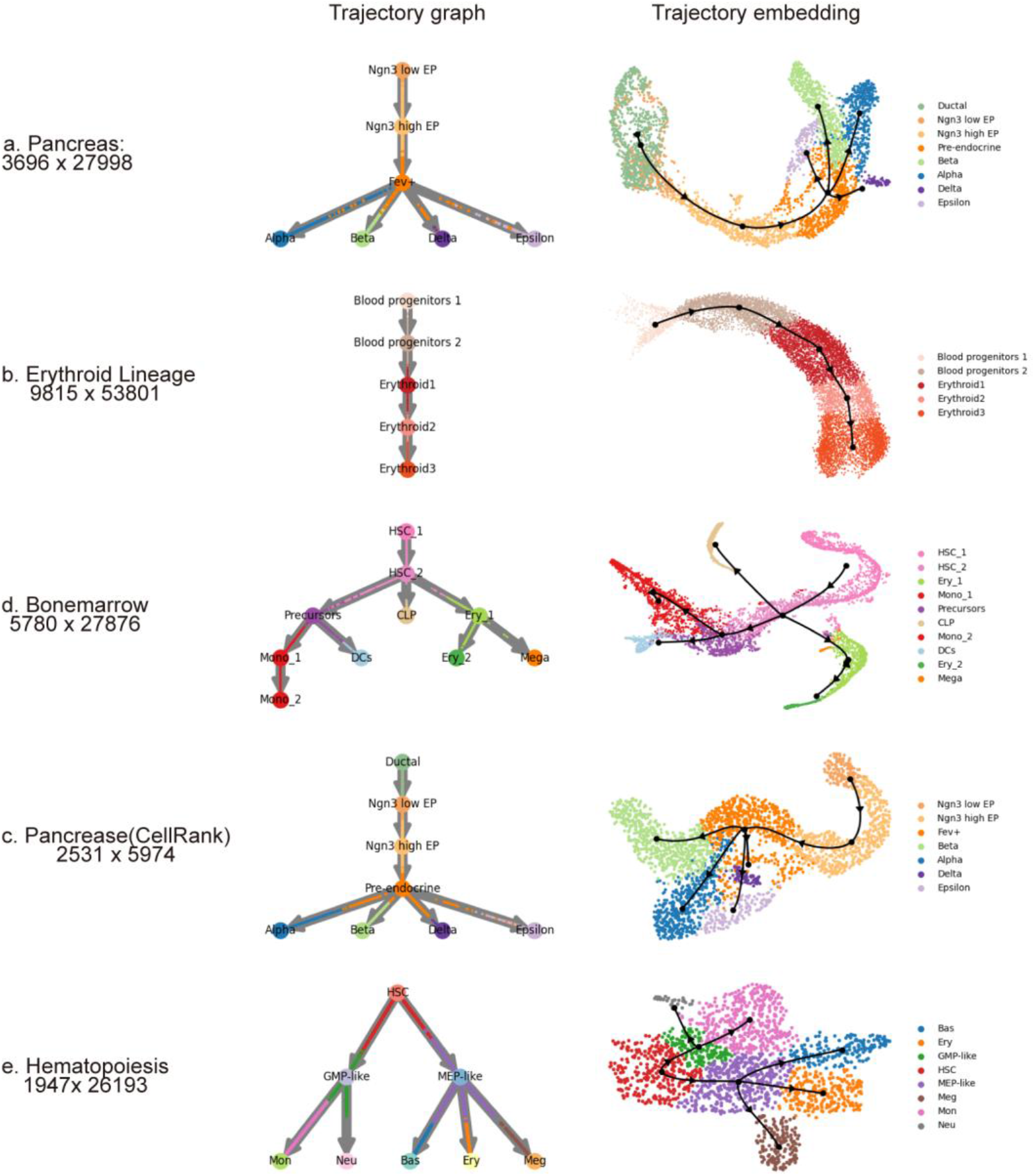
Datasets and reference trajectories preview supported by Cafe. Representative datasets bundled with Cafe are shown, including pancreas, erythroid lineage, bone marrow, CellRank pancreas, and hematopoiesis. Each row corresponds to one dataset, with the left panel showing the gold-standard trajectory graph and the right panel showing the corresponding single-cell embedding with the reference trajectory overlaid. These datasets cover diverse developmental topologies and are used to evaluate trajectory inference methods under standardized conditions.

**Extended data figure 4.**
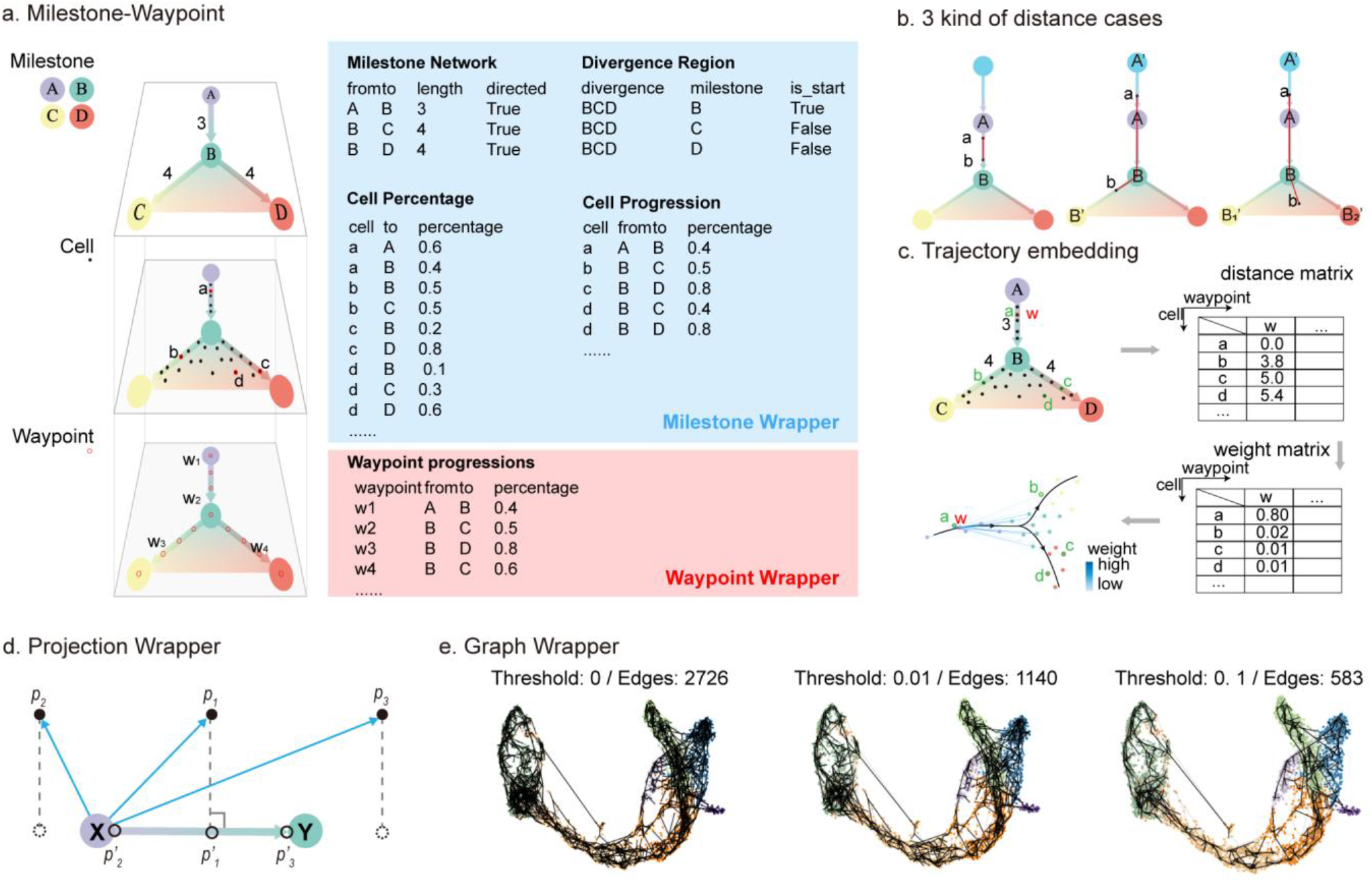
Representation of milestones, cells, and waypoints in the milestone-network framework. **a** Milestone-waypoint abstraction of the trajectory backbone, showing how the milestone network, cell percentages, divergence regions, and waypoint progressions jointly encode trajectory topology. **b**, Three representative geodesic-distance cases, including cells on the same edge, cells on different edges, and cells located in a divergence region.**c**, Waypoint embedding, in which waypoint coordinates are derived from the cell-to-waypoint distance matrix and the corresponding weights in low-dimensional space. **d**, Projection wrapper, which assigns each cell to the nearest milestone edge by geometric projection and converts its position into a progression percentage. **e**, Graph wrapper, which extracts a simplified backbone by pruning short branches at different thresholds and remaps cells onto the reduced milestone network.

**Extended data figure 5.**
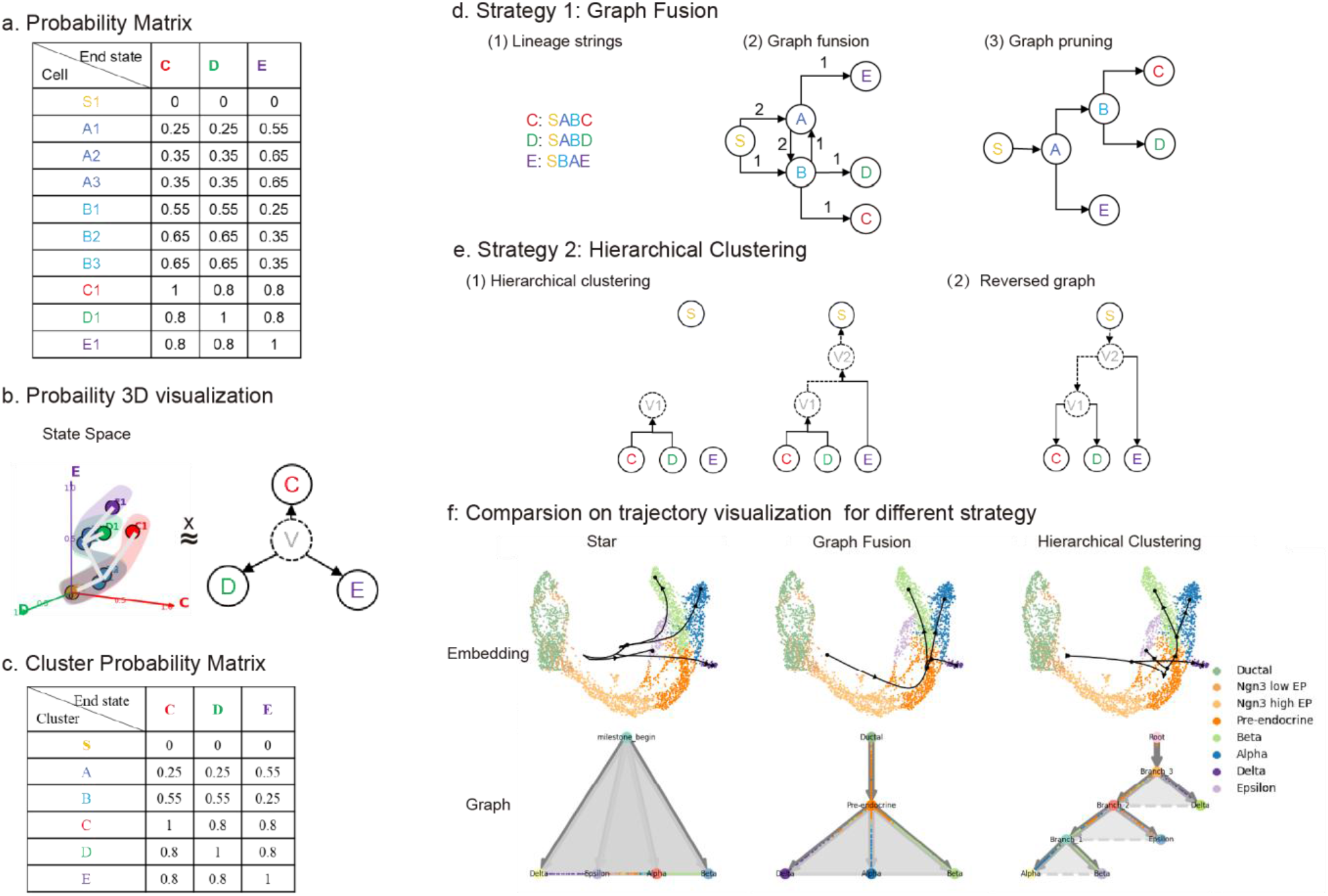
Lineage Wrapper Optimization and optimization. **a**, Probability-matrix input and the corresponding star-shaped baseline topology, in which rows represent cells and columns represent terminal states. **b**, Cluster-level probability matrix and root selection, obtained by averaging cell probabilities within clusters and identifying the cluster with the smallest total terminal-state probability as the root. **c**, Graph-fusion strategy, which orders clusters by terminal-state probability, connects adjacent states into a consensus lineage graph, and prunes redundant cycles. **d**, Hierarchical-clustering strategy, which iteratively merges terminal states according to similarity in probability space and inverts the resulting dendrogram into a rooted milestone tree.**e**, Representative comparison on the pancreas dataset, showing the star, graph-fusion, and hierarchical-clustering outputs together with their corresponding embeddings.

**Extended data figure 6.**
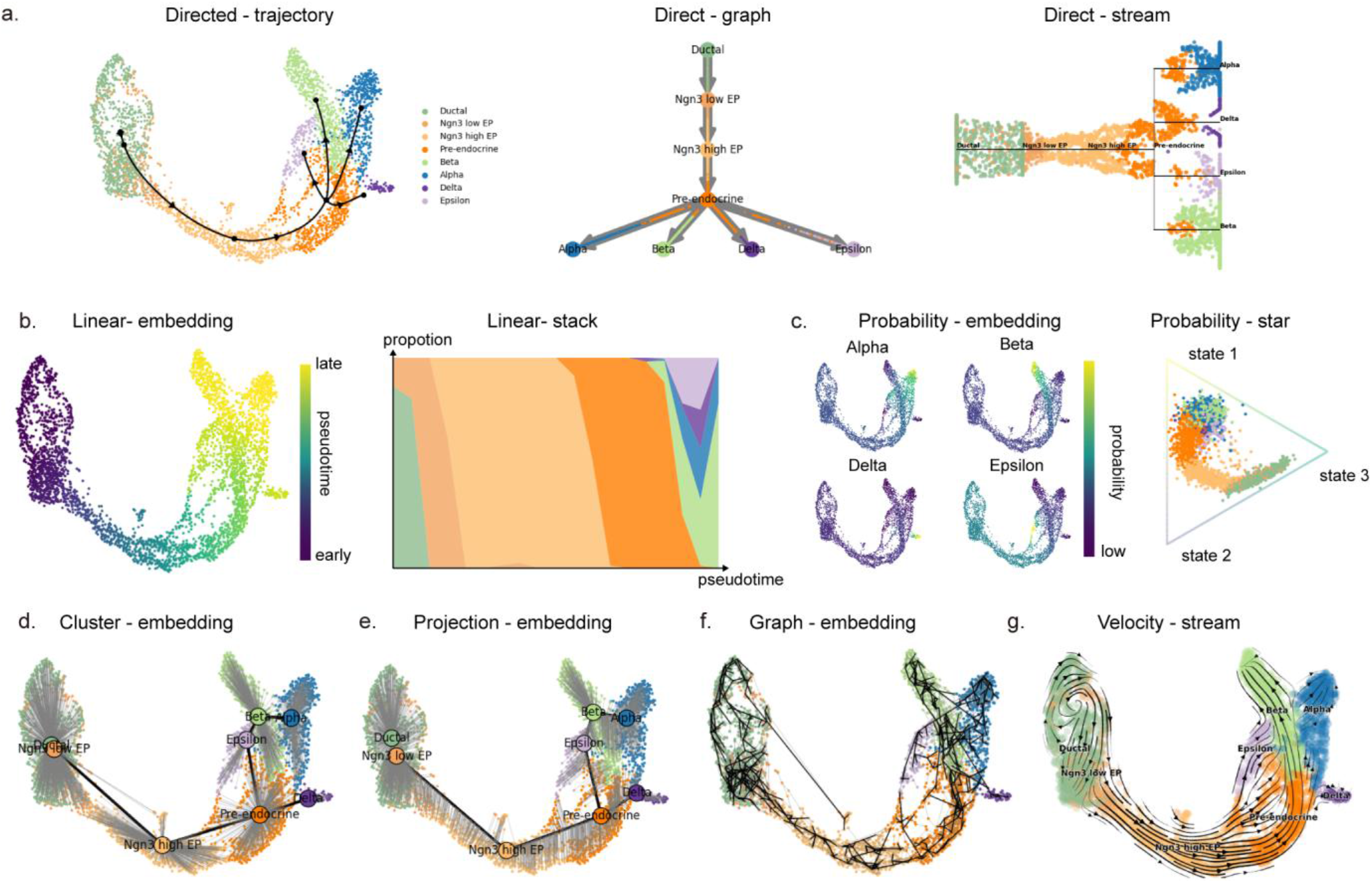
Visualization examples of wrapper outputs across representative methods. **a**, Directed-wrapper outputs, showing trajectory, graph, and stream views from a directed method.**b**, Linear-wrapper outputs, including a pseudotime-colored embedding and the corresponding stacked lineage representation. **c**, Probability-wrapper outputs, showing lineage-resolved embeddings and the corresponding star-shaped topology. **d**, Cluster-wrapper outputs, which maps cluster labels onto the inferred trajectory embedding. **e**, Projection-wrapper output, which projects cells onto milestone edges to obtain a trajectory embedding consistent with geometric assignment. **f**, Graph-wrapper output, which reconstructs backbone topology from a cell-level graph. **g**, Velocity-wrapper output, which visualizes RNA-velocity streamlines on the inferred lineage embedding.

**Extended data figure 7.**
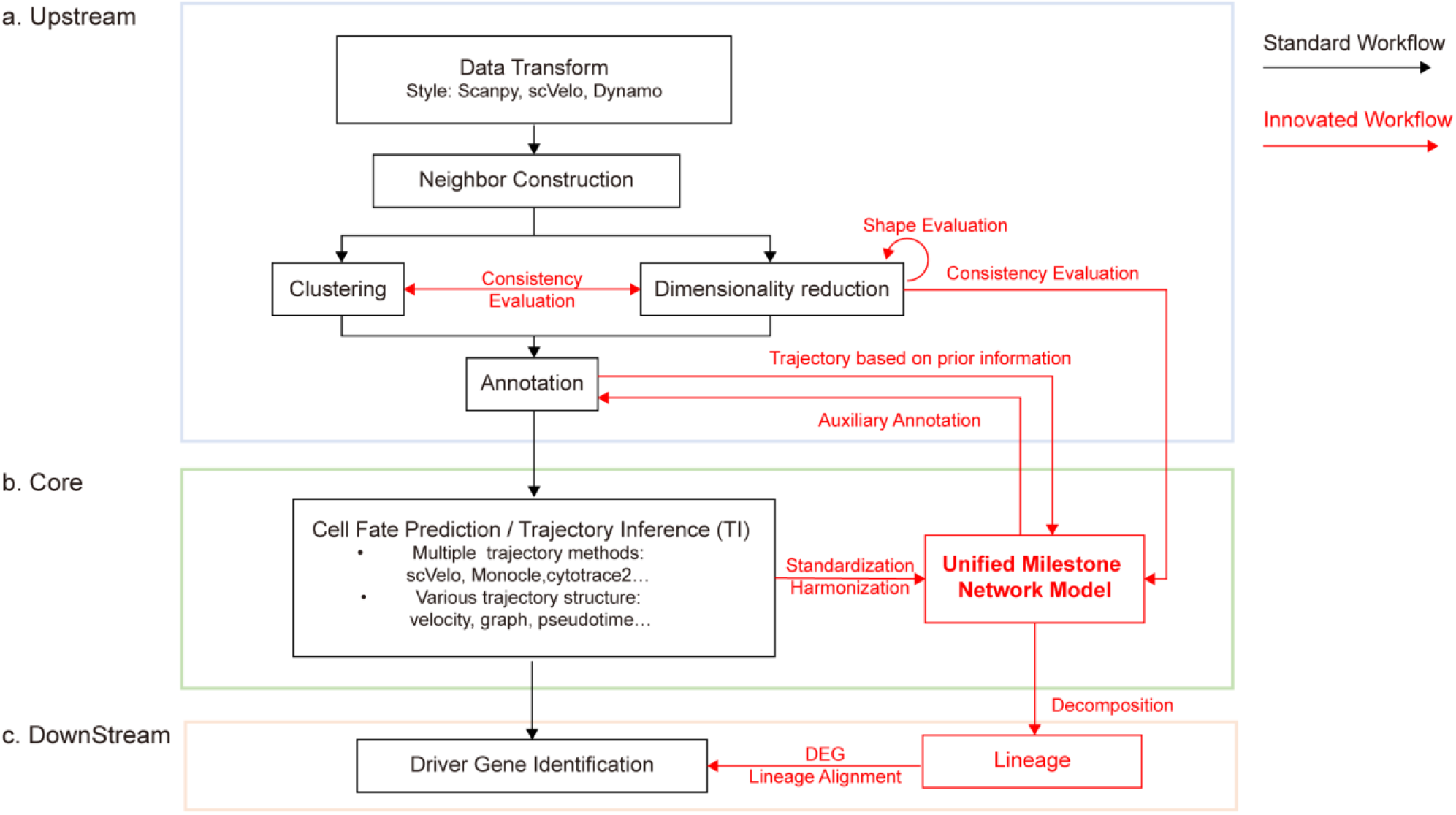
Workflow optimization of Cafe for cell fate analysis. The standard workflow is shown in black and the Cafe-optimized workflow is shown in red. The figure summarizes three stages: **a**, upstream analysis, in which trajectory-friendly embeddings are selected and annotations are incorporated when available; **b**, core analysis, in which heterogeneous trajectory inference methods are standardized into a unified milestone-network representation; **c**, downstream analysis, in which lineage-specific interpretation and driver-gene discovery are performed after decomposing the inferred topology.

**Extended data figure 8.**
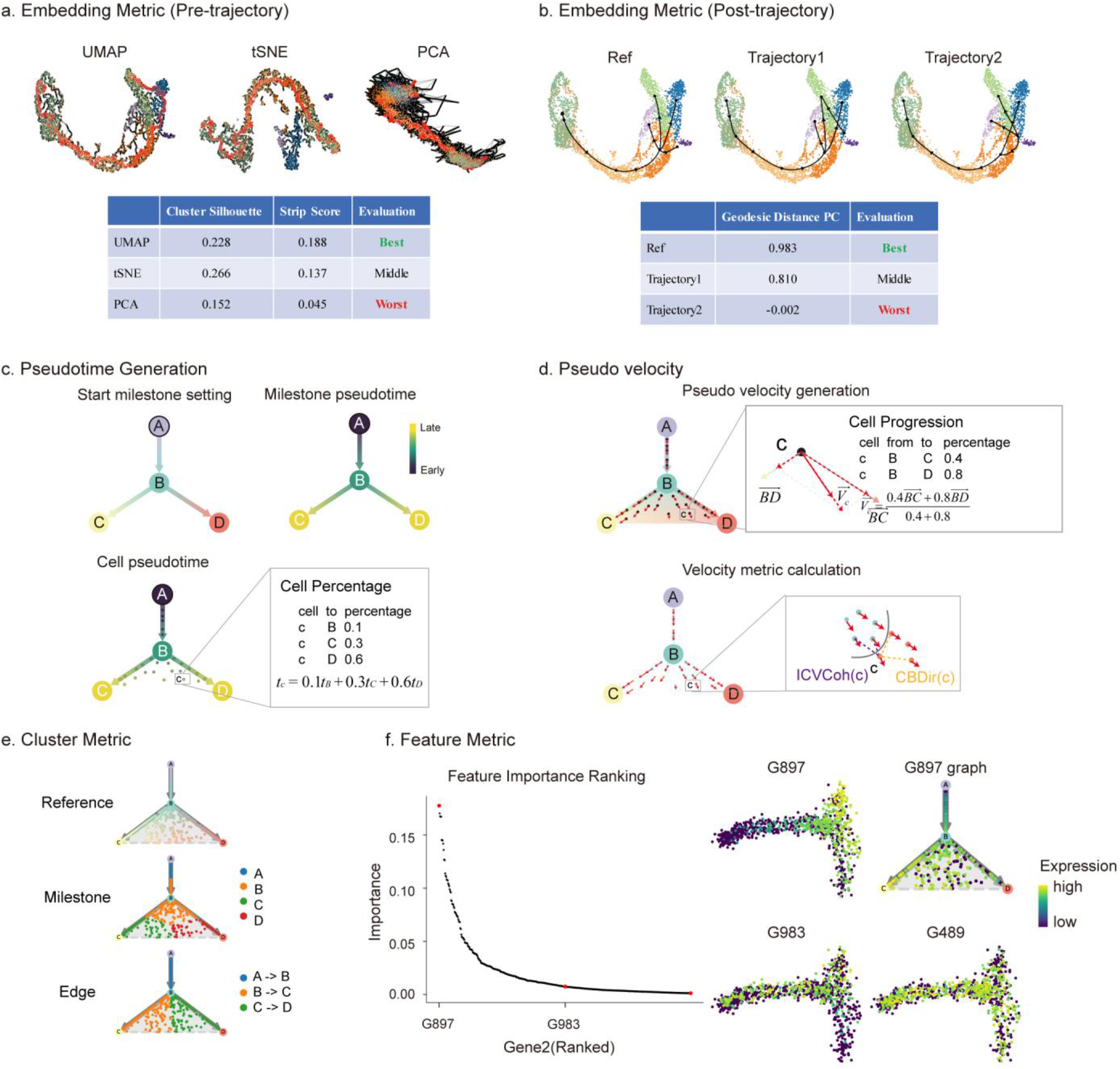
Schematic overview of metric evaluation for trajectory inference. **a**, Pre-trajectory embedding metrics, including cluster silhouette and strip score, which assess cluster separation and strip-like lineage continuity in candidate embeddings. **b**,Post-trajectory embedding metric GDPC, which measures concordance between geodesic distance in embedding space and milestone-based pseudotime. **c**, Pseudotime Correlation (PC), which quantifies agreement between predicted and reference pseudotime after graph-based or velocity-based trajectories are mapped onto a linear temporal order. **d**, Velocity metrics, including pseudo-velocity generation from milestone networks and the corresponding CBDir and ICVCoh evaluations. **e**, Cluster metric, which compares reference, milestone-based, and edge-based clustering assignments induced by the trajectory. **f**, Feature metric, where genes are ranked by feature importance and the three representative genes shown in the panel are ordered from highest to lower importance. G897 is the top-ranked candidate driver gene, showing the strongest association with trajectory progression, whereas G983 and G489 represent additional lower-ranked genes with progressively weaker feature importance and corresponding expression dynamics along the inferred lineage.

## Notes

### Competing Interest Statement

The authors have declared no competing interest.

### Summary of Updates

This version update result and method description, including framework, case study and so on.

